# Developmental hearing loss-induced perceptual deficits are rescued by cortical expression of GABA_B_ receptors

**DOI:** 10.1101/2023.01.10.523440

**Authors:** Samer Masri, Regan Fair, Todd M. Mowery, Dan H. Sanes

**Author notes:** Samer Masri, Ph.D., Center for Neural Science, New York University, 4 Washington Place New York, NY 10003. **Author Contributions**: SM, TMM, and DHS designed the experiments and wrote the paper; SM, RF, and TMM performed experiments; SM and DHS designed the viruses.

## Abstract

Even transient periods of developmental hearing loss during the developmental critical period have been linked to long-lasting deficits in auditory perception, including temporal and spectral processing, which correlate with speech perception and educational attainment. In gerbils, hearing loss-induced perceptual deficits are correlated with a reduction of both ionotropic GABA_A_ and metabotropic GABA_B_ receptor-mediated synaptic inhibition in auditory cortex, but most research on critical period plasticity has focused on GABA_A_ receptors. We developed viral vectors to express both endogenous GABA_A_ or GABA_B_ receptor subunits in auditory cortex and tested their capacity to restore perception of temporal and spectral auditory cues following critical period hearing loss in the Mongolian gerbil. HL significantly impaired perception of both temporal and spectral auditory cues. While both vectors similarly increased IPSCs in auditory cortex, only overexpression of GABA_B_ receptors improved perceptual thresholds after HL to be similar to those of animals without developmental hearing loss. These findings identify the GABA_B_ receptor as an important regulator of sensory perception in cortex and point to potential therapeutic targets for developmental sensory disorders.

**Significance Statement:** Hearing loss in children can induce deficits in aural communication that persevere even after audibility has returned to normal, suggesting permanent changes to the auditory central nervous system. In fact, a reduction in cortical synaptic inhibition has been implicated in a broad range of developmental disorders, including hearing loss. Here, we tested the hypothesis that developmental hearing loss-induced perceptual impairments in gerbils are caused by a permanent reduction of auditory cortical inhibitory synapse strength. We found that virally-mediated expression of a GABA_B_ receptor subunit in gerbil auditory cortex was able to restore two auditory perceptual skills in juvenile animals reared with hearing loss, suggesting that cortical synaptic inhibition is a plausible therapeutic target for sensory processing disorders.

## Introduction

Reduced cortical inhibition has been implicated in a broad range of developmental disorders including autism, schizophrenia, fragile x syndrome, and impaired sensory processing (Chao et al. 2010; Sanes & Kotak, 2011; Braat & Kooy, 2015; Gainey & Feldman, 2017). For example, visual or auditory deprivation that occurs during developmental sensitive periods leads to weaker inhibitory synapses between GABAergic interneurons and pyramidal cells in primary sensory cortices (Morales et al., 2022; Maffei et al. 2004; Takesian et al. 2012; Mowery et al., 2015). In some cases, these functional effects are correlated with a down-regulation of GABA receptors or loss of GABAergic terminals (Fuchs & Salazar, 1998; Kilman et al. 2002; Jiao et al. 2006; Sarro et al. 2008; Braat et al. 2015). Furthermore, when induced by hearing loss (HL), this reduction of inhibition has been linked to a broad range of perceptual and central processing deficits (Aizawa and Eggermont, 2007, Rosen et al., 2012; Yao and Sanes, 2018; Gay et al., 2014; Polley et al., 2013; Han et al., 2007; Kim and Bao, 2009; Zhang et al., 2001; Mowery et al., 2019). Taken together, these observations lead to the hypothesis that developmental HL induces a reduction of postsynaptic GABA receptor-mediated inhibition in auditory cortex (AC), thereby causing perceptual deficits. Here, we address a prediction that emerges from this hypothesis: increasing GABA_A_ or GABA_B_ receptor-dependent inhibition selectively within AC pyramidal neurons after developmental HL should restore performance on auditory perceptual tasks.

There is indirect support for the premise that normal perceptual performance is associated with appropriate levels of cortical inhibition in adults. For example, magnetic resonance spectroscopy measurements in humans demonstrate that performance on visual or auditory perceptual tasks are correlated with a higher GABA concentration (Edden et al., 2009; Dobri and Ross, 2021; Ip et al., 2021). Furthermore, a pharmacological manipulation that enhances GABAergic inhibition during behavioral testing leads to improved performance on an auditory temporal perception task in senescent gerbils and improved visual coding in senescent monkeys (Gleich et al., 2003; Leventhal et al. 2003). Consistent with this idea, systemic treatment with a GABA reuptake inhibitor can both restore the strength of inhibitory synapses following developmental HL and rescue an auditory perceptual skill (Kotak et al., 2013; Mowery et al., 2019). Although the relationship between inhibition and mature sensory processing is well established, the relative contribution of ionotropic GABA_A_ and metabotropic GABA_B_ postsynaptic receptors is uncertain. Depending on the outcome measure, pharmacological experiments suggest that both types of receptors can be an effective target for restoring normal function or plasticity (Iwai et al. 2003, Möhler et al., 2004; Fagiolini et al., 2004; Kotak et al., 2013; Cai et al., 2017; Zheng et al., 2012). Therefore, selective gain-of-function manipulations are required to determine whether restoring GABA_A_ or GABA_B_ receptor-mediated inhibition can remediate a behavioral deficit resulting from a developmental insult.

To address this problem, we developed AAV vectors to selectively increase the functional expression of either GABA_A_ or GABA_B_-mediated synaptic inhibition in the AC. One virus was designed to express the α1 subunit of the GABA_A_ receptor and the second virus was designed to express the 1b subunit of the GABA_B_ receptor, each under the CaMKII promoter (Perez-Garci et al., 2006; Vigot et al., 2006). Our approach employed a previously validated paradigm in which transient developmental hearing loss (HL) is induced during the AC critical period, causing a reduction in GABA_A_ and GABA_B_-mediated AC inhibition and diminished performance on an amplitude modulation (AM) detection task (Mowery et al., 2015; Caras and Sanes, 2015; Mowery et al., 2016; Mowery et al., 2019). We also introduce a second perceptual task, spectral modulation (SM) detection, with which to assess the effect of HL and the effect of restoring inhibition. The ability to perceive spectral modulation of sound is especially important for speech and speech-in-noise comprehension (Drullman, 1995; Zeng et al. 2005). Furthermore, humans with hearing loss or cochlear implants are impaired in spectral modulation detection, and this is correlated with speech perception (Horn et al. 2017; Ozmeral et al., 2018; Nittrouer et al., 2021). Together, AM and SM cues compose two of the fundamental building blocks of natural sounds (Singh and Theunessin, 2003), and HL-induced deficits have been linked to childhood speech and language acquisition (refs). Our findings suggest that viral expression of a GABA_B_ receptor subunit, but not a GABA_A_ receptor subunit, in the AC can remediate the deleterious effects of developmental HL on auditory perception.

## Experimental Procedures

### Experimental animals

We performed behavioral experiments on 52 normal hearing and 47 transient HL gerbils (*Meriones unguiculatus*) in the age range of postnatal days (P) 10-48. For brain slice experiments, we recorded from 34 AC pyramidal neurons, obtained from a total of 8 male and female gerbils in the age range P103-169. All animals were born from breeding pairs (Charles River Laboratories) in our colony. All procedures were approved by the Institutional Animal Care and Use Committee at New York University.

### Induction of transient hearing loss

Reversible HL was induced using earplugs made of molding clay inserted in both ears after ear canal opening, at P10, and sealed with super glue (Mowery et al., 2015; Mowery et al., 2016). Earplugs were checked daily, replaced if needed, and removed at P23. This manipulation produces a threshold shift of 15-50 dB, depending on frequency, as measured with auditory brainstem responses (Caras and Sanes, 2015), and ~25 dB at 4 kHz as measured behaviorally (Mowery et al., 2015).

### Auditory cortex brain slice recordings

Thalamocortical brain slices were generated as described previously (Kotak et al., 2005; Mowery et al., 2015, 2019). Animals were deeply anesthetized (chloral hydrate, 400 mg/kg, IP) and brains dissected into 4°C oxygenated artificial cerebrospinal fluid (ACSF, in mM: 125 NaCl, 4 KCl, 1.2 KH2PO4, 1.3 MgSO4, 24 NaHCO3, 15 glucose, 2.4 CaCl2, and 0.4 L-ascorbic acid; and bubbled with 95%O2-5%CO2 to a pH=7.4). Brains were vibratome-sectioned to obtain 300-400 μm perihorizontal auditory thalamocortical slices. The AC was identified by extracellular field responses to medial geniculate stimulation.

Whole-cell current clamp recordings were obtained (Warner PC-501A) from AC layer 2/3 pyramidal neurons at 32°C in oxygenated ACSF. Recording electrodes were fabricated from borosilicate glass (1.5 mm OD; Sutter P-97). The internal recording solution contained (in mM): 5 KCl, 127.5 K-gluconate, 10 HEPES, 2 MgCl2, 0.6 EGTA, 2 ATP, 0.3 GTP, and 5 phosphocreatine (pH 7.2 with KOH). The resistance of patch electrodes filled with internal solution was between 5-10 MΩ. Access resistance was 15-30 MΩ, and was compensated by about 70%. Recordings were digitized at 10 kHz and analyzed offline using custom Igor-based macros (IGOR, WaveMetrics, Lake Oswego, OR). All recorded neurons had a resting potential ≤-50 mV and overshooting action potentials.

Inhibitory postsynaptic potentials (IPSPs) were evoked via biphasic stimulation of layer 4 (1-10 mV, 10 s interstimulus interval) in the presence of ionotropic glutamate receptor antagonists (6,7-Dinitroquinoxaline-2,3-dione, DNQX, 20 μM; 2-amino-5-phosphonopentanoate, AP-5, 50 μM). The drugs were applied for a minimum of 8 min before recording IPSPs. Peak amplitudes of the short latency hyperpolarization (putative GABA_A_ component) and long latency hyperpolarization (putative GABAB component) were measured from each response at a holding potential (Vhold) of −50 mV. To assess GABA_B_ receptor mediated IPSPs, the GABA_A_ receptor antagonist bicuculline (10 μM) was also added to the bath. We previously verified that short- and long-latency IPSP components represented GABA_A_ and GABA_B_ receptor-dependent responses, respectively (see Fig 3D in Mowery et al., 2019).

### Behavioral training and testing

Amplitude modulation (AM) and spectral modulation (SM) depth detection thresholds were assessed with an aversive conditioning procedure (Heffner & Heffner, 1995; Kelly et al., 2006) used previously in our lab (Sarro & Sanes, 2011; Rosen et al., 2012; Buran et al., 2014; Caras and Sanes, 2015, 2017, 2019). The apparatus was controlled by custom Matlab scripts, interfaced with a digital signal processor (TDT). Stimuli were delivered via a calibrated tweeter (KEF electronics) positioned above a test cage which contains a metal water spout and floor plate. Water delivery was initiated by a syringe pump (Yale Apparatus) triggered by infrared detection of spout contact. The speaker and cage were located in a sound attenuation chamber and observed via video monitor. After placement on controlled water access, gerbils learned to drink steadily from the lick spout while in the presence of continuous, unmodulated, band-limited noise (0.1-20 kHz). Separate groups of animals were trained to withdraw from the spout when the sound changed from unmodulated noise (the “safe” cue) to either AM or SM noise (the “warn” cue) by pairing the modulation with a mild shock (0.5-1.0 mA, 300 ms; Lafayette Instruments) delivered through the spout. For the AM task, procedural training was conducted with a warn cue of 0 dB re: 100% modulation depth. For the SM task, procedural training was conducted with a warn cue of 40 dB modulation depth. Repeated pairings of the shock and the warn cue resulted in a rapidly learned association and reliable spout withdrawal, which was used as a behavioral measure of modulation detection. Warn trials were interspersed with 2-6 safe trials (2-6 seconds), during which the unmodulated sound continued unchanged; the unpredictable nature of the warn presentation prevented temporal conditioning.

After the initial associative learning, five AM or SM depths, bracketing the threshold (AM task: −3 to −27 dB re: 100% depth in 3 dB steps; SM task: +3 to 27 dB), were presented in descending order. Note that AM depth was calculated relative to a completely modulated sinusoidal waveform (dB re: 100% depth), such that larger negative values represent depths that are more difficult to detect. SM depth was calculated relative to unmodulated noise, such that smaller positive values represent depths that are more difficult to detect (dB re: 0% depth). Average stimulus level was held constant at 45 dB SPL to ensure that detection was based on the modulation cue. SM stimuli were generated using 3200 random-phase sinusoidal components spaced between 0.1 and 20kHz at a sampling rate of 44.8Khz (code courtesy of Dr. Donal Sinex). Stimuli were 1 second long with a 20 ms on and off ramps. Stimuli were generated at 2 or 10 cycles/octave. For experiments testing the effects of HL and viral vectors, SM density (2 cycles/octave) and AM rate (5 Hz) remained constant. Psychometric testing spanned 7 or 10 consecutive days. On warn trials, the response was scored as a hit when animals withdrew from the spout. On safe trials, the response was scored as a false alarm when animals incorrectly withdrew from the spout. These responses were used to calculate d’ as z(hits)-z(false alarms), a signal detection metric that accounts for individual guessing rates (Green, 1966). Values were fit with psychometric functions and used to calculate thresholds (Wichmann & Hill, 2001a, 2001b). Threshold was defined as the smallest stimulus depth at which d’ = 1.

### Development of viral vectors

Two viral vectors were developed to express either the postsynaptic GABA_B_ 1b subunit (Billinton et al., 1999), or the GABA_A_ ⍺1 subunit under the CaMKII promoter. The gene sequences for each (*Gabbr1b* and *Gabra1*) were extracted from the gerbil genome using BLAST (Zorio et al., 2019). For the *Gabbr1b* sequence, results were missing the first 141 bp of the full gene sequence, which were replaced with a sequence from the mouse to generate a complete gene sequence. These genes were inserted into viral cassettes with fluorescent reporters. At 2532 base pairs, the *Gabbr1b* sequence was too long to be used in the same cassette used for the GABA_A_ subunit while maintaining high expression levels in an AAV1 serotype. We therefore minimized cassette length by using TurboRFP as the fluorescent reporter, WPRE3 as a posttranscriptional regulatory element, and P2A to cleave the fluorescent reporter (Merzlyak et al., 2007; Choi et al., 2014, Zufferey et al., 1999; Szymczak et al., 2005). Genes were synthesized (*Gabra1*: ThermoFisher; *Gabbr1b*: Genewiz), and the full plasmids were generated, cloned, and packaged (Penn Vector Core): AAV1.CaMKII0.4.Gabbr1b.P2A.TurboRFP.WPRE3.rBG (4×10^12^ viral particles/mL), and AAV1.CaMKII0.4.Gabra1.IRES.mCherrry.WPRE.rBG (7×10^12^ viral particles/mL).

### Virus injections

For all animals, bilateral surgical virus injections into the AC were performed on P23 or P24, after earplug removal. AAV1.hSyn.eGFP.WPRE.bG (5×10^12^ vg/mL) was used as a control. Animals were anesthetized with isoflurane, and incisions made just ventral to the temporal ridge. A small burr hole (0.7 mm diameter) was drilled in the skull ~1.4 mm below the temporal ridge, just over the AC (Radtke-Schuller et al., 2016). Virus was loaded into a glass pipette backfilled with mineral oil. The glass pipettes were made sharp enough to penetrate dura without causing damage and were inserted to a depth of 400 μm. Virus was injected at a rate of 2 nl/s with a Nanoject III (Drummond), followed by a 5 min period to permit diffusion. For the two custom viruses, a volume of 600 μl was injected, whereas for the control *GFP* virus, a volume of 250 μl was injected. Incisions were closed with surgical adhesive and the animal permitted to recover for 7 days.

### Histology

At the end of each experiment, animals were deeply anesthetized with Euthasol and perfused transcardially with ice-cold PBS followed by 4% paraformaldehyde. Brains were removed and fixed in the same fixative overnight at 4°C. Brains were cryoprotected and sectioned (70 μm) on a vibratome and processed for fluorescence imaging. Only animals exhibiting TurboRFP, mCherry or GFP properly targeted to AC were included in the behavioral data.

### Statistical analysis

When data were normally distributed (as assessed by the Shapiro-Wilk W Test), values were given as mean ± SEM. Statistical analyses were conducted using MatLab. To compare multiple measures obtained from the same animal, a linear mixed effects model, correcting for subject identity (formula: threshold ~ 1 + group * training day + (1 | subject)) was used to verify a main effect of treatment group or the interaction between training day and treatment group. For individual training days, group comparisons of variables were made using a one-way analysis of variance (ANOVA) followed by least significant differences post-hoc. The HL + GFP group was specified as the control group for all post hoc tests. The best threshold achieved by each animal during 7 or 10 days of psychometric testing (depending on the experiment) was used to determine perceptual thresholds. The significance level was set at α = 5%. All data are expressed as mean ± SEM unless otherwise stated.

## Results

To investigate the causal relationship between AC synaptic inhibition and HL-induced perceptual deficits, we developed viral vectors to express the gerbil gene sequences for subunit α1 of the GABA_A_ receptor (*Gabra1*) or subunit 1b of the GABA_B_ receptor (*Gabbr1b*), each under a CAMKII promoter (see Methods). Our reasoning was that overexpression of a single subunit of either receptor would remove a rate limiting step in the surface expression of the functional multimeric receptor, thereby increasing IPSP amplitude.

### Functional assessment of virally expressed GABA_A_ and GABA_B_ receptor subunits

To determine whether each viral vector led to functional upregulation of cortical inhibitory postsynaptic potentials (IPSPs), we performed whole cell current clamp recordings from pyramidal neurons in auditory cortex brain slices 21-27 days after virus injection into AC (Figure 1A; see Methods). IPSPs were elicited in response to local electrical stimulation, as described previously (Mowery et al., 2016; Mowery et al., 2019). Figure 1B shows the vector used to express the *Gabra1* subunit under a CaMKII promoter, and a fluorescence image of the reporter molecule, mCherry, as observed during whole cell recordings. Figure 1C shows representative IPSPs recorded from a *Gabra1*-infected (orange trace; yellow pipet in panel B) and an uninfected AC neuron (gray trace) at a holding potential (V_hold_) of −50 mV in the presence of glutamate receptor blockers (20 μM DNQX; 50 μM AP-5). Figure 1D shows that the peak amplitudes of the short latency IPSP hyperpolarization (putative GABA_A_ component, labeled “A”) was significantly greater for infected neurons (Mean ± SEM; infected: 12.8 ± 0.6 mV; uninfected: 6.8 ± 0.6 mV; q=2.05, df=26, p<0.0001). Therefore, the *Gabra1* vector increased GABA_A_ receptor-mediated IPSP amplitude in AC pyramidal neurons.

**Figure 1:**
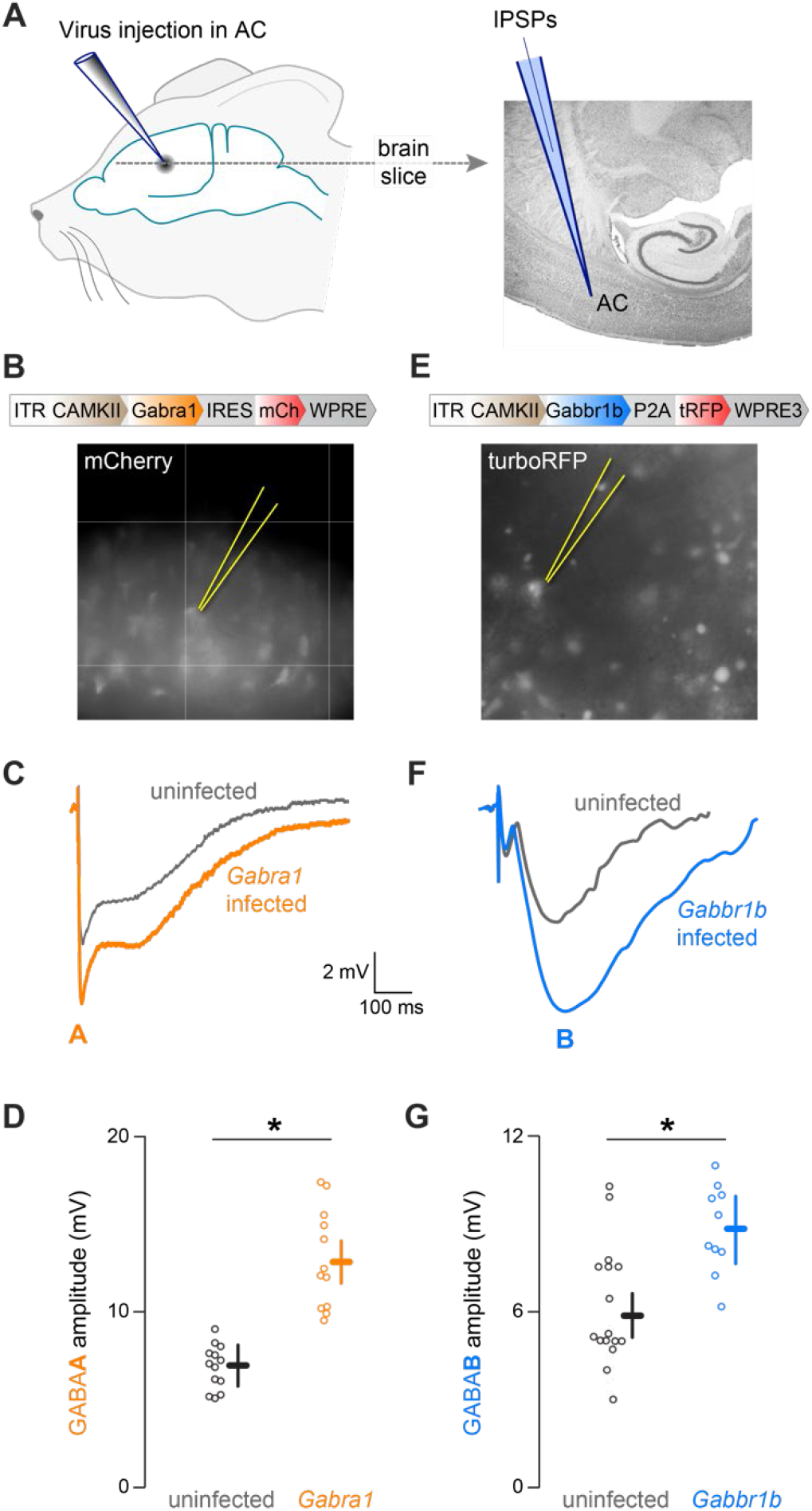
Viral vector design and validation. (A) For both Gabra1 and Gabbr1b AAVs, primary auditory cortex layer 2/3 was injected (Nanoject 2; Drummond) with approximately 250 nl of virus. After three weeks of expression a thalamocortical slice preparation was made and whole cell recordings (current clamp) from ACx L2/3 pyramidal cells were carried out. (B) Top, Diagram showing Gabra1 vector. Bottom, micrograph from ACx L2/3 showing Gabra1 infected cells (fluorescing, mCh) and one patched pyramidal neuron. (C) Representative evoked IPSP showing the larger GABAA potential in the Gabra1 infected pyramidal neuron (fluorescing patched cell from B) vs local uninfected (non-fluorescing) pyramidal neuron from the same slice. D) Plot diagram showing the average difference in GABA_A_ IPSP amplitudes for uninfected versus Gabra1 infected pyramidal neurons. (E) Top, Diagram showing Gabbr1b vector. Bottom, micrograph from ACx L2/3 showing Gabbr1b infected cells (fluorescent, mCh) and one patched pyramidal neuron. (F) Representative evoked IPSP showing the larger GABA_B_ potential in the Gabbr1b infected pyramidal neuron (fluorescing patched cell from C) vs local uninfected (non-fluorescing) pyramidal neuron from the same slice. (G) Plot diagram showing the average difference in GABA_B_ IPSP amplitudes for uninfected versus Gabbr1b infected pyramidal neurons.

Figure 1E shows the vector used to express the postsynaptic *Gabbr1b* subunit (Billinton et al., 1999) under a CamKII promoter, and a fluorescence image of the reporter molecule, turboRFP, as observed during whole cell recordings. Figure 1F shows representative IPSPs recorded from a *Gabbr1b*-infected (blue trace; yellow pipet in panel E) and an uninfected AC neuron (gray trace) at a holding potential (V_hold_) of −50 mV in the presence of glutamate receptor antagonists (20 μM DNQX; 50 μM AP-5) and a GABA_A_ receptor antagonist (10 μM bicuculline). Figure 1G shows that the peak amplitudes of the long latency IPSP hyperpolarization (putative GABA_B_ component, labeled “B”) was significantly greater for infected neurons (Mean ± SEM; infected: 8.7 ± 0.6 mV; uninfected: 5.8 ± 0.4 mV; q=2.03, df=32, p=0.0002). Therefore, the Gabbr1b vector increased GABA_B_ receptor-mediated IPSP amplitude in AC pyramidal neurons.

#### Assessing perceptual performance on AM and SM tasks

Figure 2 outlines the full experimental protocol. We induced reversible developmental hearing loss, beginning at ear canal opening (P10) and ending after the auditory critical period (P23) by inserting earplugs (Figure 2A, orange shading), as described previously (Caras and Sanes, 2015). We injected separate groups of HL-reared animals with the *Gabra1* (n = 8) or *Gabbr1b*-expressing vector (n = 9), or a GFP control virus (n = 9), bilaterally in AC between P23 and P25, after earplug removal. We also included a normal hearing group (n = 15; Figure 2A, gray shading). Following a 7 day recovery period, animals were water restricted and began behavioral training on P30.

**Figure 2:**
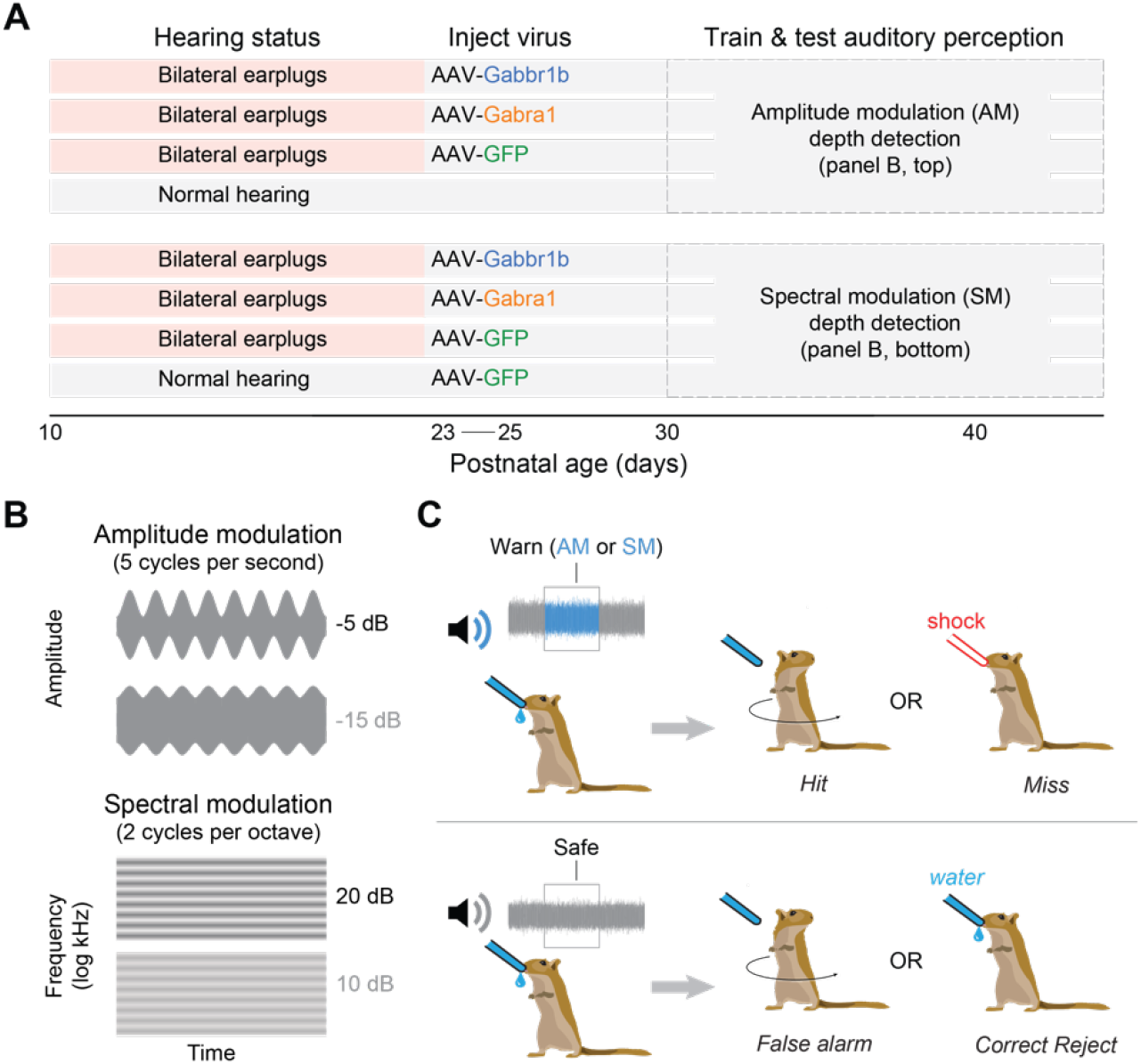
Experimental paradigm. (A) The experimental timeline, and each of the experimental groups is shown. (B) Example stimulus waveforms are shown for the AM depth detection task (top) and the SM depth detection task (bottom). (C) The Go-Nogo paradigm used for psychometric testing is shown.

Separate groups of animals were tested on AM depth detection (Sarro and Sanes, 2011; Rosen et al., 2012; Caras & Sanes, 2015) or SM depth detection. As described in Methods, control and HL-reared animals were trained to drink from a lick spout during continuous noise (0.1-20 kHz, 45 dB SPL). For the AM detection task, animals were initially trained to withdraw from the spout when 5 Hz amplitude modulation at 0 dB re: 100% occurred (Figure 2B, top). For the SM detection task, animals were initially trained to withdraw from the spout when 2 cycles/octave density spectral modulation at 40 dB depth occurred (Figure 2B, bottom). Procedural learning continued until animals achieved a d’ ≳ 1.3 with a 0 dB re: 100% depth AM stimuli or 40 dB depth SM stimuli (4-8 days). We then conducted 7 days of psychometric testing as animals’ performance gradually improved on the AM or SM task (Figure 2C). Perceptual thresholds improved as gerbils responded to smaller modulation depths due to perceptual learning (Caras & Sanes, 2017).

We first tested the effect of HL and GABA receptor subunit expression on AM depth detection. Figure 3A presents representative psychometric functions for two individual animals and shows that AM detection was superior for the HL-reared gerbil that received bilateral AC injections of a *Gabbr1b*-expressing vector (HL+*Gabbr1b*; blue line) as compared to the HL-reared gerbil that received AC bilateral injections of a *GFP*-expressing vector (HL+*GFP*; green line). As schematized in Figure 2A, top, we obtained thresholds for animals in these two groups, as well as HL-reared animals that received a *Gabra1*-expressing vector (HL+*Gabra1*), and normal hearing (NH) animals.

**Figure 3:**
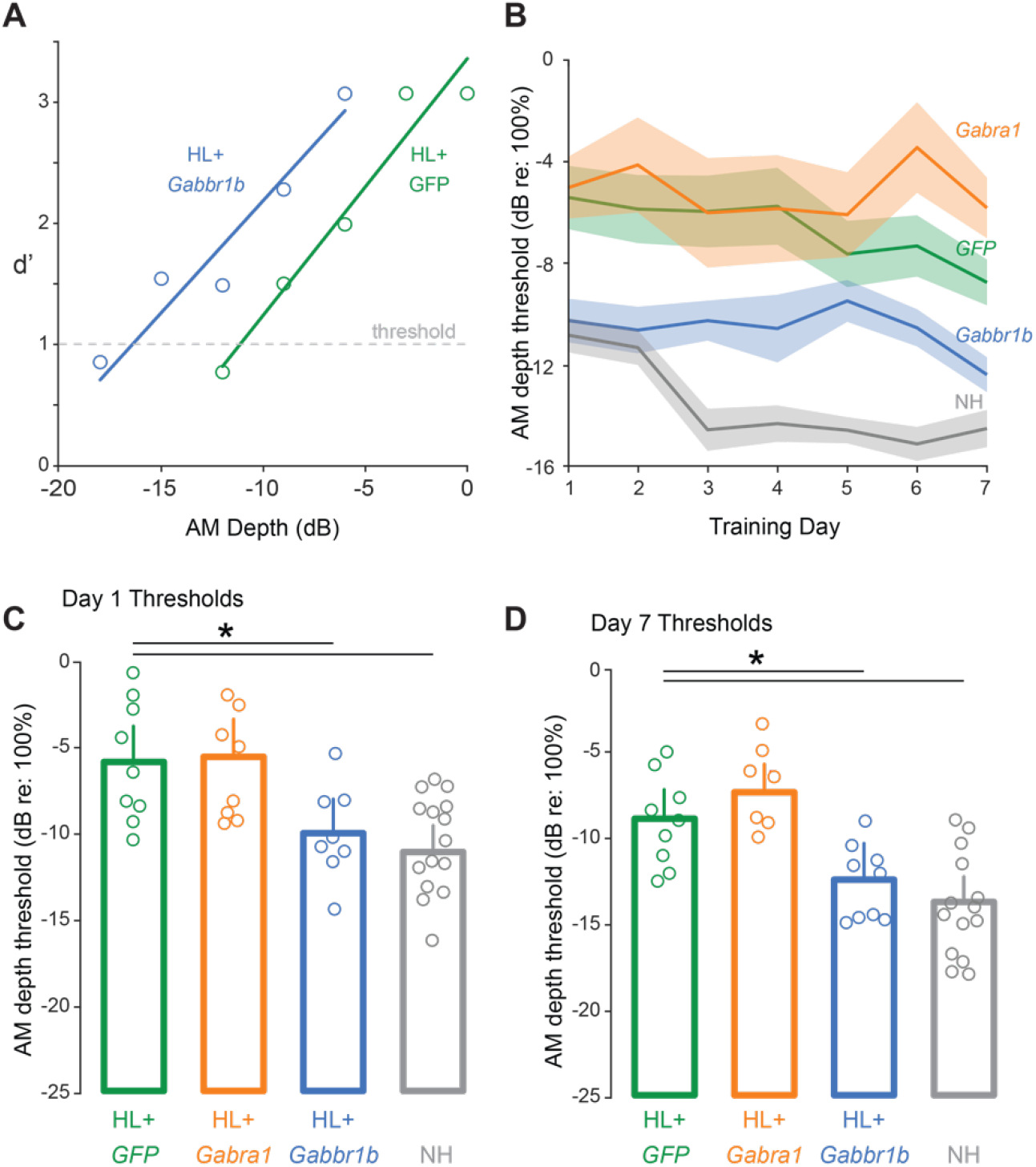
Gabbr1b expression restores AM detection. (A) Representative behavior for a HL-reared gerbil expressing *Gabbr1b* (HL+*Gabbr1b*) and a HL-reared gerbil expressing *GFP* (HL+*GFP*) in AC, both tested after transient hearing loss (HL). (B) AM depth thresholds achieved by each group over training days. Mean ± SEM. (C) *Gabbr1b* expression in AC rescued AM perception relative to *GFP* expression on day 1 of psychometric testing. Bars indicate significant differences (see text for statistical values). (D) *Gabbr1b* expression in AC rescued AM perception relative to *GFP* expression on day 7 of psychometric testing. Bars indicate significant differences (see text for statistical values).

Figure 3B shows thresholds by group over 7 days of psychometric testing. A linear mixed-effects model comparing the effects of training day and virus condition on thresholds, and taking into account individual subject behavior, indicates that viral treatment was a significant factor (F = 10.83, p = 9.83 x 10^−7^). Therefore, expression of the *Gabbr1b* subunit restored normal behavioral performance on the AM detection task in HL-reared animals in a manner that was independent of training day.

There was a significant effect of treatment group on day 1 of perceptual testing (one-way ANOVA, p = 4.02 x 10^−5^, F = 10.46, df = 40, Figure 3C) and day 7 of perceptual testing (one-way ANOVA, p = 6.85 x 10^−8^, F = 20.07, df = 40, Figure 3D). A post hoc comparison revealed that AM detection thresholds were significantly poorer for transient HL-reared animals that received a control virus (HL+*GFP*) as compared to normal hearing animals (NH). This was the case both for day 1 (HL+*GFP* = −5.61 ± 0.99 dB, NH = −10.94 ± 0.77 dB, p = 8.35 x 10^−4^) and day 7 of testing (HL+*GFP* = −8.90 ± 0.91 dB, NH = −14.56 ± 0.70 dB, p = 1.00 x 10^−4^). This finding confirms the effect of HL reported previously (Caras and Sanes, 2015; Mowery et al., 2019).

Post hoc comparisons also revealed that GABA_B_ subunit expression, but not GABA_A_ receptor expression could partially restore AM detection thresholds in HL-reared animals. Animals in the HL+*Gabbr1b* group displayed significantly lower AM detection thresholds than the HL+*GFP* group, both at day 1 (HL+*Gabbr1b* = −10.37 ± 0.99 dB, p = 0.009, NH = −10.94 ± 0.77 dB, p = 8.35 x 10^−4^) and day 7 of perceptual testing (HL+*Gabbr1b* = −12.47 ± 0.91 dB, p = 0.04). In contrast, the HL+*Gabra1* group did not different significantly from the HL+*GFP* group, either at day 1 (HL+*GFP* = −5.61 ± 0.99 dB, HL+*Gabra1* = −5.24 ± 1.06 dB, p = .99), or day 7 of perceptual testing (HL+*GFP* = −8.90 ± 0.91 dB, HL+*Gabra1* = −6.01 ± 0.96 dB, p = .15).

### SM detection performance in normal hearing gerbils

To develop a behavioral test of perception of spectral modulation (SM), we modified the aversive conditioning paradigm such that unmodulated white noise transitioned to spectrally modulated noise on “warn” trials. Similar to the AM detection task, SM stimuli were presented at multiple depths in 3 dB increments to determine each animal’s perceptual threshold. Since SM detection has not been assessed previously in gerbils, we first sought to validate two features of this psychometric task. SM is described by a density (cycles/octave) which specifies the peak-to-trough distance of the logarithmic sinusoidal frequency filter used. Humans achieve their best thresholds in the range of 2-4 cycles/octave and thresholds increase at 10 cycles/octave (Eddins and Bero, 2007). We trained gerbils on the same aversive conditioning paradigm used for AM, but with the change cue being a SM stimulus at either 2 or 10 cycles/octave. We found that best thresholds achieved during 10 days of psychometric testing were significantly lower at 2 cycles/octave (5.4 ± 1.43 dB, n = 14) than 10 (8.3 ± 2.5 dB; p = 0.363, t = −2.148, df = 11, unpaired t-test, n = 8; Figure S1). Therefore, all subsequent SM detection psychometric tests used 2 cycles/octave. To confirm that gerbil SM detection is robust to changes in level, as reported fo humans (Eddins and Bero, 2007), we alternated between 45 dB SPL and 36 dB SPL during 4 additional days of testing with 2 cycles/octave stimuli (n = 8). There was no significant difference in thresholds over each pair of testing days (45 dB days: 9.2 ± 4.0 dB, 36 dB days: 10.66 ± 5.03 dB, p = 0.562, t = 0.593, df = 14, paired t-test). This indicates that gerbil perception of SM stimuli is similar to that displayed by humans.

### *Gabbr1b* expression restores SM detection following developmental hearing loss

We next tested the effect of HL on SM depth detection, and the ability of GABA receptor expression to rescue a HL-induced perceptual deficit. Figure 4A presents representative psychometric functions for two individual animals and shows that SM detection was superior for the HL-reared gerbil that received bilateral AC injections of a *Gabbr1b*-expressing vector (HL+*Gabbr1b*; blue line) as compared to the HL-reared gerbil that received AC bilateral injections of a *GFP*-expressing vector (HL+*GFP*; green line). As schematized in Figure 2A, bottom, we obtained thresholds for animals in these two groups, as well as HL-reared animals that received a *Gabra1*-expressing vector (HL+*Gabra1*), and normal hearing (NH) animals (HL+*GFP*, n = 8; HL+*Gabra1*, n = 7; HL+*Gabbr1b*, n = 6; NH+*GFP*, n = 7). As observed previously with procedural training on the AM detection task, all 4 groups reached criterion in a similar number of trials and reached similar maximum d’ on the SM detection task (Figure S2).

**Figure 4:**
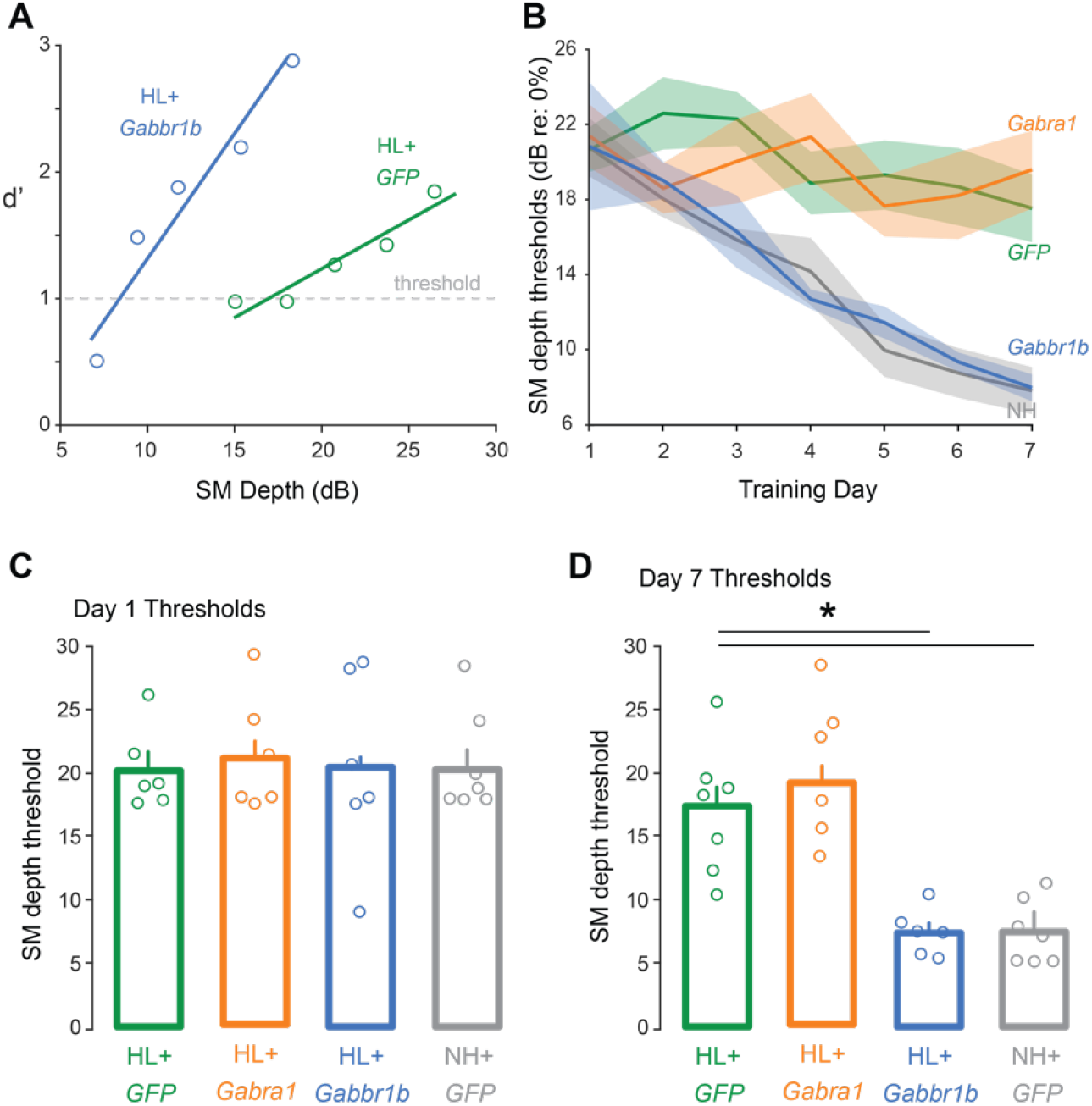
Gabbr1b expression restores SM detection. (A) Example psychometric curves of individual gerbils showing d’ at each of the 5 modulation depths presented in a single session. The leftward shift of the HL+*Gabbr1b* function, relative to HL+*GFP* function indicates improved performance. Bars indicate significant differences (see text for statistical values). (B) Group performance on each day of psychometric testing. (C) There are no differences in SM modulation thresholds on the first day of psychometric testing, as calculated by fit crossing d’ = 1. (D) SM thresholds on day 7 of psychometric testing. Both HL+*Gabbr1b* and NH+*GFP* groups performed significantly better than HL+*GFP* animals. Bars indicate significant differences (see text for statistical values).

Figure 4B shows group thresholds over all 7 days of training. A linear mixed effects model shows that viral treatment alone is not a significant factor (F = 0.634, p = 0.594) but the interaction between group and training day was (F = 13.257, p = 7.64 x 10^−8^). Therefore, expression of the *Gabbr1b* subunit restores normal threshold improvement trends on the SM detection task in HL-reared animals.

The SM detection thresholds of all four groups did not differ significantly from one another on day 1 of psychometric testing (HL+*GFP*: dB = 20.66 ± 2.11; HL+*Gabra1*: dB = 21.38 ± 2.11; HL+*Gabbr1b*: dB = 20.83 ± 2.31; NH+GFP: dB = 20.77 ± 1.96; ANOVA, p = .9951, F = 0.02, df = 23, Figure 4C). However, as shown in Figure 4D, a significant effect of treatment group emerged by day 7 of testing (one-way ANOVA, p = 1.34 x 10^−5^, F = 14.5, df = 27). NH+*GFP* and HL+*Gabbr1b* animals both reached low thresholds of 7.8 ± 1.6 dB and 8.0 ± 1.8 dB, respectively. In contrast, thresholds for HL+*GFP* (17.5 ± 1.5 dB) and HL+*Gabra1* (19.6 ± 1.6 dB) animals improved very little over 7 days of testing. There was no significant difference between day 7 thresholds for HL+*Gabbr1b* and NH+*GFP* animals and both were significantly better than HL+*GFP* (p = 0.002, p = 0.0011, respectively).

## Discussion

Proper regulation of synaptic inhibition is integral to the development and maintenance of sensory processing. Transient or permanent developmental HL that begins during an AC critical period causes a long-lasting reduction of cortical synaptic inhibition that is attributable to the functional loss of both ionotropic GABA_A_ and metabotropic GABA_B_ receptors (Kotak et al., 2005; Takesian et al., 2012; Mowery et al., 2019). This reduction of AC inhibition correlates with impairments in psychometric performance on a range of auditory tasks as well as degraded AC neuron stimulus processing (Rosen et al., 2012; Buran et al., 2014; Caras and Sanes, 2015; Ihlefeld et al., 2016; von Trapp et al., 2017; Yao & Sanes, 2018). To test whether there is a causal relationship between AC inhibition and perceptual skills, we upregulated GABA receptor-mediated inhibition in AC pyramidal neurons through viral expression of the *Gabra1* or *Gabbr1b* subunit genes in animals reared with HL. Our results show that upregulating GABA_B_ receptor-dependent inhibition through expression of the gerbil *Gabbr1b* subunit gene can rescue two different perceptual deficits, AM and SM detection. In contrast, upregulating GABA_A_ receptor-mediated inhibition through expression of the gerbil *Gabra1* subunit gene had no effect on perceptual performance. Therefore, our results suggest that the magnitude of AC inhibition is positively correlated with perceptual performance, with postsynaptic GABA_B_ receptors playing a pivotal role.

Amplitude and spectral modulation are discrete features of natural sounds, including speech, and sensitivity to these cues is correlated with speech comprehension (Cazals et al., 1994; Shannon et al., 1995; Singh and Theunissen, 2003; Elliott 2009; Nittrouer et al., 2021). Here we developed a behavioral paradigm to assess SM detection in gerbils, such that the effects of HL could be compared with a previously characterized percept, AM detection (Rosen et al., 2012; Caras & Sanes, 2015). We found that rearing conditions and viral treatment had no effect on procedural training times, in agreement with past results with our AM paradigm, although training times were comparatively longer (Figure 2S). Adult humans display depth detection thresholds of ~2 dB at 2 cycles/octave, with poorer performance at higher spectral densities (Eddins and Bero, 2007). In agreement, we found that gerbils have better thresholds at 2 cycles/octave than 10 cycles/octave and that best thresholds are within 3 dB of human performance at 2 cycles/octave (Figure 1S). We also analyzed a concatenated series of hundreds of gerbil vocalizations and found that spectral modulation drops off significantly above 2 cycles/octave (not shown). This validates our use of a new perceptual test with which to assess the impact of hearing loss and the capacity of treatments to restore perception.

### Interpreting the pattern of restored AM and SM detection following *Gabbr1b* expression

*Gabbr1b* expression rescued AM and SM detection in HL-reared animals, but with different magnitudes and time courses. For the AM detection task *Gabbr1b*-treatment improved perceptual performance in HL-reared animals from the first day of testing as compared to GFP-treated controls (Figure 3). This improved performance was maintained during the 7 days of testing. This outcome is consistent with a rapid improvement in AM stimulus encoding following *Gabbr1b* expression, but no effect on perceptual learning (i.e., an improvement in detection threshold as a result of practice). In contrast, *Gabbr1b* expression led to a gradual improvement of SM thresholds during the 7 days of testing, identical to NH animals with GFP expression (Figure 4). In principle, these differences in the effect of restoring inhibition could relate to differences in the way that AM and SM stimuli are represented in the AC.

Amplitude and spectrally modulated noise are expected to differ in terms of the evoked discharge pattern of AC neurons. AM stimuli are known to produce a strong temporal response that correlates with the AM rate. In contrast, SM stimuli produce a response that is dependent on frequency tuning (e.g., inhibitory sidebands) (Calhoun & Schreiner, 1994, 1998; Atencio & Schreiner, 2010). At the cellular level, one possibility is that feedforward inhibition mediated by Parvalbumin-expressing interneurons tightens the timing of the auditory evoked response in the input Layer 4/5 (Wehr and Zador, 2003; Nocon et al., 2022), which may contribute to perception of amplitude modulation. In contrast, SM stimuli are stationary. Here, intracortical pathways may recruit local interneurons that mediate lateral inhibition, increasing gain in Layer 2/3, thereby improving the detection of energy differences across spectral bands (Kaur et al., 2004; Kaur et al., 2005; Li et. al. 2014).

### Relationship of GABA receptor manipulation to the AC critical period

In the current experiments, the manipulations and behavioral assays all occur prior to sexual maturation, a time during which inhibitory functional properties continue to mature (Pinto et al., 2010; Takesian et al., 2012). A large body of research from the developing visual pathway shows that inhibitory synapse development regulates cortical plasticity. Monocular deprivation (MD) leads to reduced cortical activation by the deprived eye during a developmental CP, and experimentally increasing GABAergic transmission can close the critical period prematurely (reviews: Hensch, 2004; Hensch, 2005; Hooks and Chen, 2007). One implication of these observations is that inhibition in adult animals is too strong to permit plasticity. In fact, manipulations that reduce cortical inhibition in adults can induce excitatory synaptic plasticity (He et al., 2006; Sale et al., 2007; Fernandez et al., 2007; Harauzov et al., 2010). Therefore, when inhibitory strength is high, behavioral deficits can be ameliorated by temporarily lowering it. In contrast, our results suggest that when inhibitory strength is low, as occurs after developmental HL, behavioral deficits can be ameliorated by permanently raising it.

The gerbil AC displays a well characterized critical period (CP) for the effect of HL that closes at P18 (Mowery et al., 2015). When HL is initiated after P18, there is no reduction to AC inhibitory synapse strength (Mowery et al., 2016). In contrast, when HL is initiated before P18, the reduction of AC inhibitory synapse strength persists to adulthood and can be attributed to the functional loss of both GABA_A_ and GABA_B_ receptor-mediated IPSPs (Mowery et al., 2019). Therefore, a core premise of this study is that the loss of one or both forms of postsynaptic inhibition is causally related to perceptual deficits that attend developmental HL. Since the virus was injected into AC on P23, our results suggest that the manipulation need not occur during the cortical CP in order to restore normal neural and behavioral function. This is consistent with our finding that systemic treatment with a GABA reuptake inhibitor (SGRI) from P23-35 also rescues AC inhibition and AM detection thresholds (see Fig 2e in Mowery et al., 2019).

Since a reduction of GABA_A_ receptor mediated inhibition has been implicated in many developmental disorders, it was reasonable to predict that upregulating inhibitory strength through *Gabra1* expression (Figure 1D) would rescue HL-induced deficits on auditory tasks. However, we previously reported that systemic treatment with a GABA_A_ ⍺1 receptor agonist, zolpidem, does not restore AM detection thresholds following development HL (see Fig 2f in Mowery et al., 2019). Two lines of evidence may explain why perceptual performance was rescued only by upregulating GABA_B_ receptor-mediated inhibition. First, GABA_B_ receptor function may have a direct impact on synaptic plasticity, particularly during development. Postsynaptic GABA_B_ receptors can induce inhibitory long-term depression (iLTD) at feedforward inhibitory synapses between Parvalbumin-expressing interneurons and Pyramidal neurons in input layers of visual cortex during a developmental CP (Wang and Maffei, 2014). This mechanism has been implicated in auditory map remodeling (Vickers et al., 2018), and GABA_B_ receptor agonists enhance ocular dominance plasticity (Cai et al., 2017). Second, postsynaptic GABA_B_ receptors are located extrasynaptically and modulate both the activity of postsynaptic GABA_A_ receptors and NMDA receptor-driven LTP (Komatsu, 1996; Fritschy et al., 1999; Charara et al., 2005; Booker et al., 2013; Tao et al., 2013, Connelly et al. 2013). GABA_B_ receptor activation may also induce BDNF release, thereby inducing the addition of perisomatic GABAergic synapses (Fiorentino et al., 2009). Therefore, although both forms of GABAergic inhibition are reduced by HL, dysregulation of GABA_B_ receptor’s modulatory role may have a substantial impact on the acquisition of perceptual skills during development.

### Limitations to data interpretations

Expression of AAV vectors begins within one week of infusion, and typically ramps up to a plateau at 2-3 weeks (Reimsnider et al. 2007; Kaplitt et al. 1994; Chamberlin et al. 1998). Therefore, we timed our experiment such that psychometric testing fell within a 2-3 week post-injection window during which time expression should have been maximal. However, it is possible that a continued increase of expression during training was associated with a greater influence on behavioral performance at later testing days. A second consideration is that this study used two specific inhibitory receptors, but the approach did not restrict expression to synapses from any particular type of inhibitory interneuron. Parvalbumin or Somatostatin-expressing interneurons at the thalamorecipient layer or in L2/3, which naturally target GABA_B_ receptors each synapse onto pyramidal neurons that expressed GABBR1B protein following viral infection (Manz et al. 2019). Additionally, L2/3 Pyramidal neurons extend dendrites to L1 within which Neurogliaform interneurons form synapses that primarily evoke GABA_B_ receptor-mediated responses and regulate plasticity (Tamás et al., 2003). Therefore, specific interneuronal connections responsible for the reported behavioral effects is unknown. Finally, it is possible that increasing inhibitory transmission could have induced homeostatic upregulation of excitation in AC and a maintenance of excitatory/inhibitory balance (Turrigiano & Nelson, 2004; Le Roux *et al*., 2006). This potential effect could include a normalization of excitatory cellular properties and may have contributed to the behavioral benefits of our manipulation. In fact, GABA_A_ receptors have been shown to regulate homeostatic plasticity and this may explain why expressing GABRA1 did not improve performance (Wen et al. 2022; Le Roux et al. 2008; Rannals & Kapur, 2011).

### Conclusion

Here we have shown that restoring inhibition in AC alone was sufficient to restore auditory perception after a developmental insult. Restoring cortical synaptic inhibition may be relevant to a range of developmental disorders. For example, GABA levels in visual cortex are reduced in amblyopia and this is correlated with weaker perceptual suppression by the amblyopic eye (Mukerji et al., 2022). By directly comparing the impact of restoring *Gabra1* and *Gabbr1b* protein expression we have shown that this effect was only achieved by upregulating GABA_B_ receptor-mediated inhibition. This result is surprising given a long focus on GABA_A_ receptor-mediated inhibition in the field of developmental sensory processing (Fagiolini et al. 2004; Chang et al., 2005). Our findings suggest that regulating GABA_B_ receptor mediated inhibition remains a plausible target for therapies that seek to prevent or reverse behavioral deficits that attend developmental disorders.

## Abbreviations

AC: auditory cortex
*Gabra1*: gamma-aminobutyric acid A receptor subunit ⍺1
*Gabbr1b*: gamma-aminobutyric acid B receptor subunit 1b
HL: hearing loss
IPSP: inhibitory postsynaptic potential
AM: amplitude modulation
SM: spectral modulation

## Acknowledgements

The work is supported by NIDCD R01DC011284 (DHS and TMM), and F32DC020659 (SM).

---

**Supplemental Figure 1:**
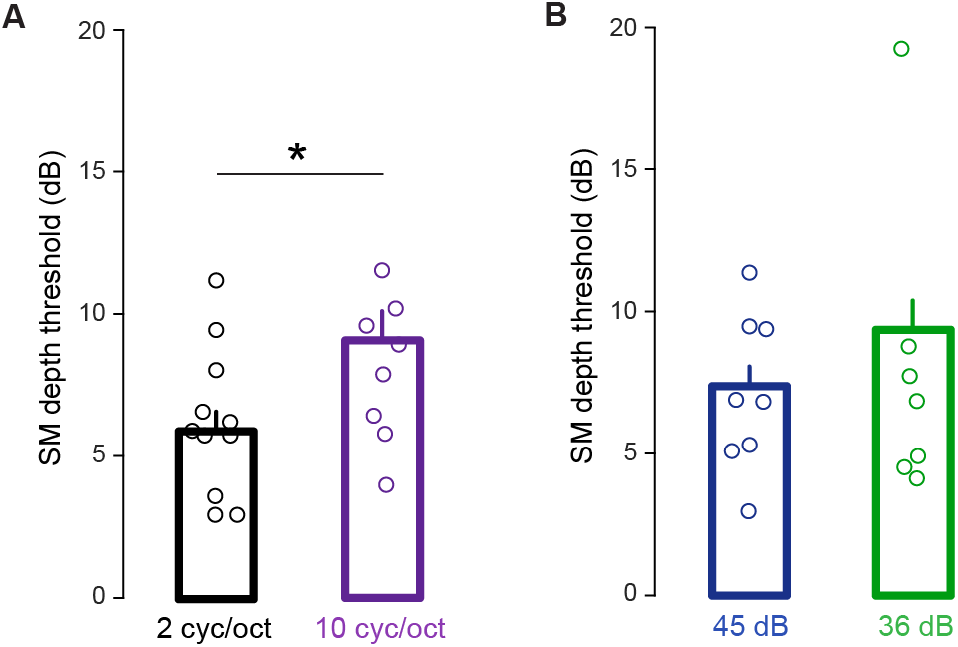
Spectral modulation detection in normal hearing juvenile gerbils. (A) NH animals display better thresholds for SM at 2 cycles/octave relative to 10. Bar indicates significant difference (see text for statistical value). (B) SM detection thresholds do not change significantly when sound is presented at a lower level of 36 dB SPL (p = 0.593).

**Supplemental Figure 2:**
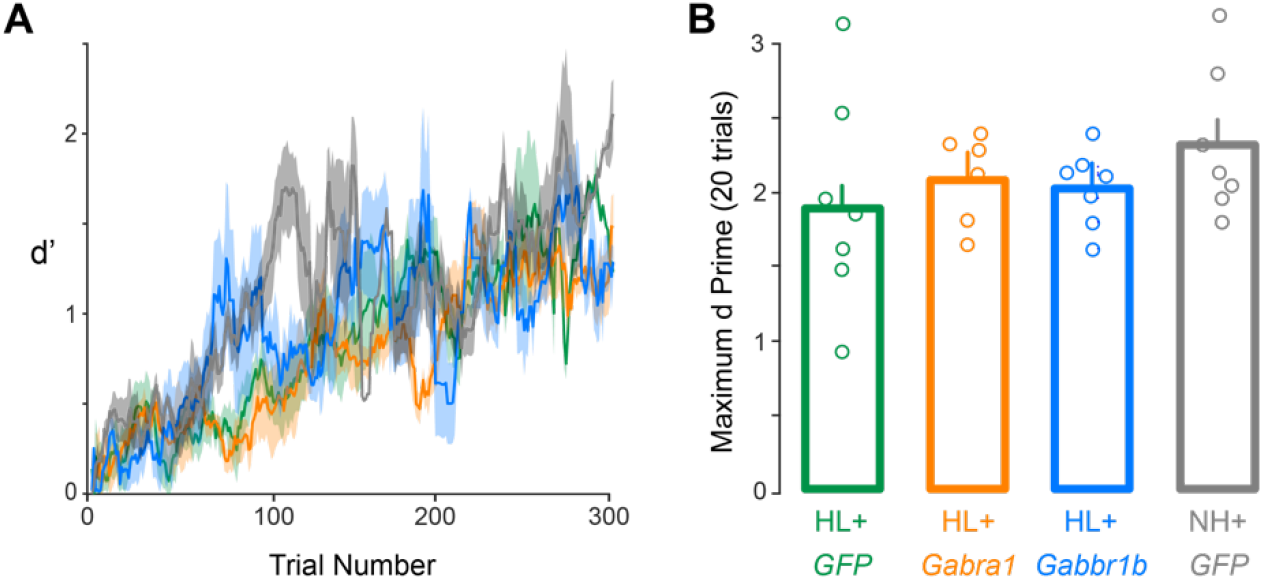
Procedural training for spectral modulation detection. (A) There are no group differences in the number of trials required for procedural training (mean ± SEM, moving window of 20 trials). (B) There are no significant differences in maximum d’ achieved during all procedural training over 20 trial windows (see text for statistical value).

